# Drug screening targeting TREM2-TYROBP transmembrane binding

**DOI:** 10.1101/2024.12.02.626344

**Authors:** M. Cobas-Carreño, A. Esteban-Martos, L. Tomas-Gallardo, I. Iribarren, L. Gonzalez-Palma, A. Rivera-Ramos, J. Elena-Guerra, E. Alarcon-Martin, R. Ruiz, M.J. Bravo, J.L. Venero, X. Morató, A. Ruiz, J.L. Royo

## Abstract

*TREM2* encodes a microglial membrane receptor involved in the disease-associated microglia (DAM) phenotype whose activation requires the transmembrane interaction with TYROBP. Mutations in *TREM2* represent a high-impact risk factor for Alzheimer’s disease (AD) which turned TREM2 into a significant drug target. We present a bacterial two-hybrid (B2H) system designed for high-throughput screening of modulators for the TREM2-TYROBP transmembrane interaction. In a pilot study, 315 FDA-approved drugs were analyzed to identify potential binding modifiers. Our pipeline includes multiple filtering steps to ensure candidate specificity. The screening suggested two potential candidates that were finally assayed in the human microglial cell line HMC3. Upon stimulation with anti-TREM2 mAb, pSYK/SYK ratios were calculated in the presence of the candidates. As a result, we found that varenicline, a smoking cessation medication, can be considered as a transmembrane agonist of the TREM2-TYROBP interaction.

## Introduction

TREM2 is a transmembrane receptor expressed in most innate immune cells including microglia. The expression of TREM2 in these cells underscores its key role in maintaining homeostasis in the brain and other tissues, and in regulating inflammatory and repair processes. Microglial TREM2-dependent pathway favors the structural integrity and functionality of brain synapses, protecting from the cognitive decline associated to Alzheimer’s disease (AD) (Leng and Edison, 2021; Fracassi et al., 2023). Different studies suggested that TREM2 could exert an anti-inflammatory action and increased activation of the TREM2 pathway could be associated to resilience against the harmful effect of Aβ plaques (Fracassi et al., 2023). Under this scenario, AL002c was launched, consisting in a mouse IgG1 anti-hTREM2 monoclonal antibody (mAb) generated using the recombinant hTREM2 extracellular domain as an immunogen in a hybridoma approach, followed by humanization and affinity maturation by yeast display (Wang S, et al 2020). Shortly after, a humanized version of the monoclonal antibody, was released which counteracts TREM2 decreased functionality by optimizing its signaling to improve cell survival and proliferation, and activity of microglia and is currently under examination in a Phase II clinical trial (Clinical trials.gov/INVOKE-2). In parallel, other strategies focus on the use of small molecules rather than mAb. It has been recently initiated the Phase I clinical trial of VG-3927, a novel small molecule TREM2 agonist, to treat common neurodegenerative diseases associated with microglia. However, evidence suggests that activated microglia could contribute to neuronal death in advanced stages of the disease suggesting that the TREM2-dependent pathway might act as a double-edged sword in the AD (Konishi H, et al 2018; Long et al., 2019; Qin et al., 2021; Garcia-Alberca JM et al 2024). With the objective of increasing our pharmacological arsenal against this therapeutic target, we have developed a high throughput screening strategy focused on the identification of potential modulators of the TREM2-TYROBP interaction. More specifically, we have focused our target to their transmembrane binding, in an attempt to find molecules with high hydrophobicity scores that eventually would facilitate the blood-brain barrier penentrance.

## Materials and methods

### Plasmids and strains

Constructs contained in the N-terminal site the *Pseudomonas aeruginosa* phage Pf3 coat protein signal peptide (QSVITDVTGQLTAVQADITTIGG) (Kiefer D, et al 1999) followed by the transmembrane domain (TMD) of the either TYROBP or TREM2 and a cytoplasmic 3-glycine elbow to increase flexibility in order to facilitate the interaction between the two *Bordetella pertussis* adenylate cyclase complementary domains (T18 and T25) of the BATCH dual hybrid system (Battesti A et al 2012). The bacterial two-hybrid (B2H) system based on adenylate cyclase which generates cAMP what in the *Escherichia coli* BTH101 background turns into increasing beta-galactosidase expression. Plasmids containing the chimeric fusions were ordered from Genscript (https://www.genscript.com/) and further subcloned using conventional molecular biology techniques in DH5α *E. Coli* strain. The complete protein used as bait is translated from pKNT25 of the in-frame cloned BATCH pKTN25 plasmid (Supplementary material)

**Table 1.**

| **Strains** |  |
| --- | --- |
| ***E. coli* DH5α** | fhuA2 lac(del)U169 phoA glnV44 Φ80' lacZ(del)M15 gyrA96 recA1 relA1 endA1 thi-1 hsdR17. |
| ***E. coli* BTH101** | Str<sup>R</sup>. lacZ<sup>+</sup> strain that responds to intracellular cAMP levels of the BACTH system. Genotype: F<sup>-</sup>; cya-99, araD139, galE15, galK16, rpsL1 (Str), hsdR2, mcrA1, mcrB1. |
| **Plasmids** |  |
| **pKNT25-TYROBPTM** | Km<sup>R</sup>. Plasmid with the transmembrane domain of human TYROBP fused to the NH<sub>2</sub>- terminus of the T25 fragment of adenylate cyclase together with the signal peptide Pf3. First component of the initial drug screening system. |
| **pUT18-TREM2TM** | Ap<sup>R</sup>. Plasmid with the transmembrane domain of human TREM2 fused to the NH<sub>2</sub>- terminus of the T18 fragment of adenylate cyclase together with the signal peptide Pf3. Second component of the initial drug screening system. |
| **pKNT25-zip** | Km<sup>R</sup>. Positive control plasmid (ZIP) for the BACTH system. First component of the system for discarding non-specific inhibitors. |
| **pUT18-zip** | Ap<sup>R</sup>. Positive control plasmid (ZIP) for the BACTH system. Second component of the system for discarding non-specific inhibitors. |
| **pKNT25** | Km<sup>R</sup>. Plasmid backbone prepared to clone at the amino terminus of the T25 fragment of the adenylate cyclase of the BACTH system. First component of the nonspecific activator discard system. |
| **pUT18** | Ap. Plasmid backbone prepared to clone at the amino terminus of the T18 fragment of the adenylate cyclase of the BACTH system. Second component of the nonspecific activator discard system. |

### Cell culture assays

HMC3 cells (ATCC, Manassas, VA, USA) were maintained in DMEM medium supplemented with 10% fetal bovine serum, GlutaMAX, and 1% penicillin/streptomycin/fungizone (Thermo Fisher Scientific, Waltham, MA, United States). Semiconfluent (50-70%) cultures were stimulated with 1-7 μg/ml anti-TREM2 (1:1 anti-TREM2 R&D systems AF1729; anti-TREM2 R&D MAB17291) for 5 minutes at 37ºC in the presence of 1% dimetilsulfoxide (DMSO) alone or in combination with the candidate drug. Plates were then rapidly placed on ice and washed twice with phosphate buffer saline, *in situ* lysated with TRIsure and stored O/N at -80ºC for at least 24h. Proteins were fractionated by electrophoresis using 10% sodium dodecyl sulphate (SDS) polyacrylamide gels, electroblotted into PVDF membranes (Hybond-P, GE Healthcare), and blocked with 5% BCA in TBS. Membranes were then incubated with the different antibodies overnight at 4 °C (anti-hSYK #2710, anti-hPhospho-SYK #2712 from Cell Signaling were used according to manufacturer’s instructions), followed by incubation with a horseradish peroxidase-conjugated antibody. The immunoreactive bands were visualized using ECL (Invitrogen). The images were obtained and analyzed with ChemiDoc™ Touch Imaging System (Bio-Rad).

### Semiquantitative bacterial two hybrid

Prior to each assay, plasmids were freshly transformed in *E. coli* BTH101 and plated at 37 ° C overnight (O/N). Four colonies were selected from each plate, added separately to 1 mL of LB with 100 ng/µL ampicillin and 50 ng/µL of kanamycin, and stirred at 1200 rpm overnight at 37° C. The next day, 10 µL drops of each biological replica were seeded in triplicate on plates with both antibiotics and grown at 30°C. After 24h, a picture was taken to analyze the intensity of blue of each drop with the ImageJ software (W. Rasband, National Institutes of Health). The blue color correlates with the amount of cAMP generated, linked to the interaction capacity of the study proteins, and was normalized with respect to the negative control (BTH101 with pUT18 and pKNT25). For the analysis, a 32-bit image type was selected, the different rows were defined with the Analyze Gel command, then the baseline was defined with Plot Lanes and the interior of the peak area was chosen to obtain the result.

### Quantitative bacterial two hybrid

Freshly transformed *E. coli* BTH101 with the necessary plasmids were grown O/N at 37°C in 0.5 mL of LB with both ampicillin 100 μg/ml and kanamycin 20 μg/ml. 1:50 dilutions were grown in 10 mL tubes at 30°C and 180 rpm until OD_600_ reached 0.2-0.3. Culture was then divided in 200 μl aliquots in flat-bottom 96-well plates where drugs (FDA-Approved Drug Library HY-L022, MedChem Express, Monmouth Junction, USA) were assayed in triplicates at 150 μM. Positive controls contained 150 ng/mL IPTG and DMSO 1%. Plates were sealed to avoid evaporation and incubated O/N at 180 rpm and 30ºC. For the enzymatic assay 100 µL of diluted 1:10 aliquots were taken and OD_620_ was determined in an ELISA reader. 50 µL of these diluted samples were added to 705 µL of Z buffer (60 mM Na_2_HPO_4_ 7H_2_O; 40 mM NaH_2_PO4 H_2_O; 10 mM KCl; 1 mM MgSO_4_ 7H_2_O) with 0.3% of freshly added ß-mercaptoethanol, 30 µL of chloroform and 15 µL of 0.1% SDS. Samples were vortexed for 10 sec and stabilized for 2 min in a water bath at 30°C. The reaction started adding 200 µL ONPG (4 mg / mL in Z buffer) and stopped with 0.5 ml Na_2_CO_3_ 1M before being placed on ice. Samples were then spined and placed in an ELISA 96-well plate for OD_405_ measurement. ELISA reader filters available were 620 nm and 405 nm instead of the 600 and 420 defining the Miller Units. Correlations between the 600 and 420nm absorbances taken with a conventional spectrophotometer and the 620 and 405 nm with the ELISA plate reader were R^2^>0.98, (p-value <0.001, Spearman’s test) and allowed us to define the Adapted Miller Units (UMAs) as: 1000×[OD_405_ / (t × 0,05 × OD_620_)] (Supplementary Figure 1).

### Molecular Docking

The 3D structure for Varenicline (DB01273) was obtained from the Drugbank Olie database (Knox C et al 2024) and the TREM2 and TYROBP α-helices were extracted from our previous molecular dynamics simulation (Garcia-Aberca JM et al, 2024). The input files for the docking studies for the helices and the ligand were prepared using the AutoDockTools (Morris GM, et al, 2009) software package and the docking simulation was performed using the AutoDock Vina (Eberhadt J et al 2021; Trott O et al 2021) software. The box was set to cover all the inter-helices space (size_x, _y and _z = 36, 32, 86 and center_x, _y, _z = 49.696, 46.055, 32.111) and the energy_range and exhaustiveness parameters were set to 4 kcal/mol and 8 respectively. The results of the docking analysis were visualised using PyMOL software (The PyMOL Molecular Graphics System, Version 1.2r3pre, Schrödinger, LLC, Germany). The NCI establish between the ligand and the two helices were analysed using the NCIPLOT4 software package and visualised using Jmol with a density isososurface of 0.4 and a colour range of [-2,2] with blue representing attractive interactions, red representing repulsive interactions and green representing the weaker interactions.

## Results

Drop assays of *E. coli* BTH101 strain harboring different plasmid configurations showed that both human TMDs effectively bind in the bacterial membrane (Mean of 1.4 vs 1.01, p-value<0.01, Mann Whitney test), as predicted by dynamic models and according to what previously reported (Figure 1). However, the dynamic range of this beta-galactosidase assay was lower than expected. For this reason we decided to perform the screening in liquid cultures.

**Figure 1.**
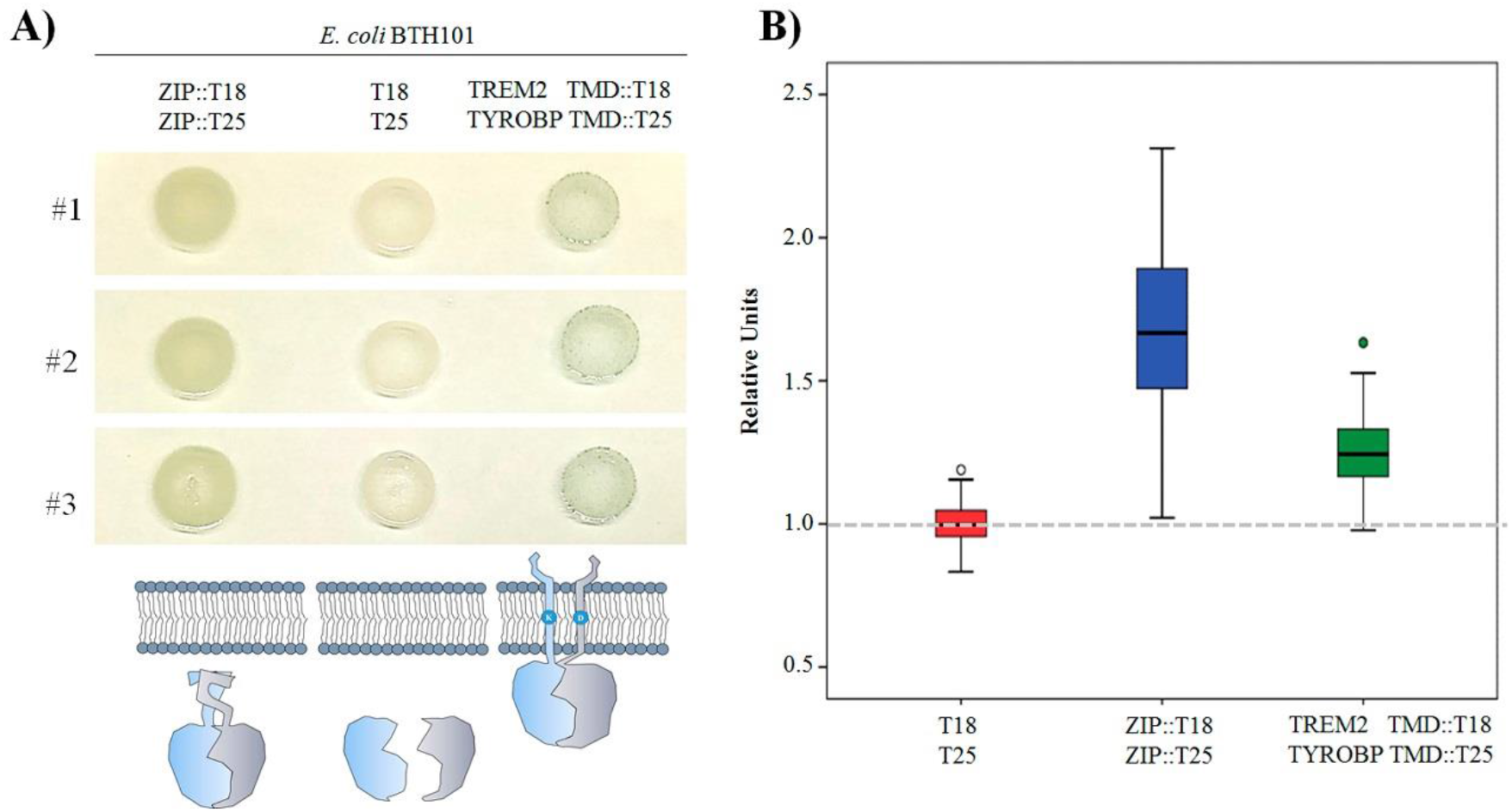
Transmembrane BATCH two hybrid system. A) Three independent drops *of E. coli* BTH101 strain harboring different plasmid configurations grown overnight at 30ºC in LB X-gal plates with Km and Ap. ZIP domains allowed a cytoplasmic reconstitution of the adenylate cyclase as the TREM2 and TYROBP TMD do in the membrane turning blue the drops. Color background obtained from the T18 and T25 control plasmids comes from fortuitous cytoplasmic binding. B) Box plot showing median, quartiles and range values of the relative intensities compared to the empty plasmids. Semiquantitative analysis of the X-gal drops under the different configurations (T18/T25 n=99; ZIP::T18/ZIP::T25 n=81; TREM2TMD::T18/TYROBP TMD::T25 n=57).

Thus, we tested 315 FDA-approved drugs which were assayed in triplicate and compared with the negative control (Figure 2A). We selected the most promising candidates 38 drugs: 25 potential activators and 13 repressors, which represented 8% and 4% of the assayed compounds, respectively. These were individually assayed to perform an independent assay to validat the modulating capacity of 13 of these drugs (10 activators and 3 inhibitors). This bias in the identification of activators (3.2%) vs inhibitors (1%) was statistically significant (p=0.049, χ^2^ test) (Supplementary table 1).

**Figure 2.**
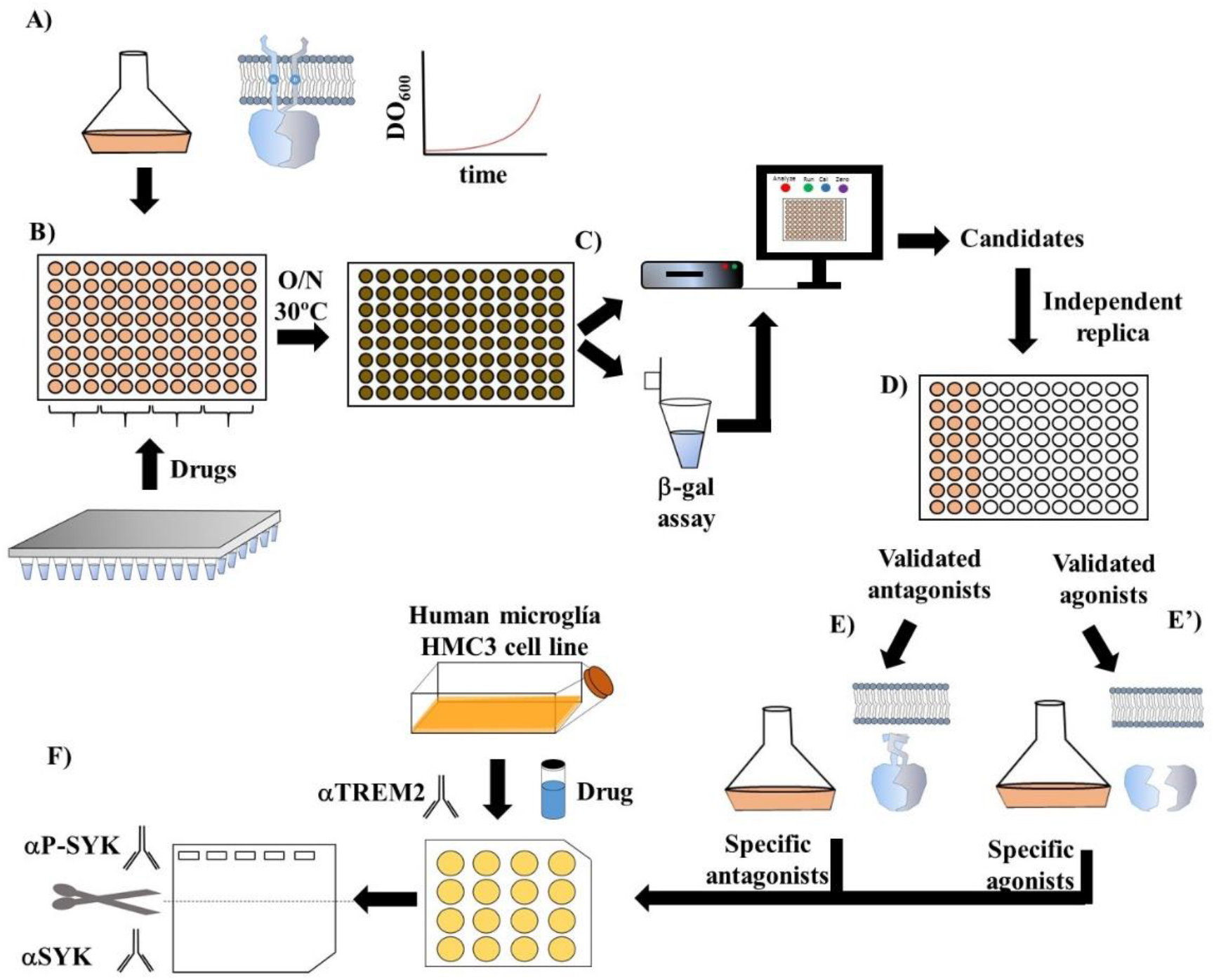
Screening pipeline. A) *E. coli* BTH101 bearing the two plasmids with TREM2TMD::T18 and the TYROBPTMD::T25 was cultured until OD_600_=0-2-0.25. B) Exponential-phase bacteria were distributed into a 96-well plate in combination with the different drugs. Plates were incubated o/n at 30ºC. C) Cells were diluted 1:10 and growth was calculated as OD at 620 nm. Beta-galactosidase assay were performed and OD_405_ was also calculated. D) Candidates were independently repeated resulting in validated agonists and antagonists. E) To determine specificity, true antagonist drugs were challenged to BTH101 stain bearing ZIP::T18 and ZIP::T25. E’) similarly, true agonist were challenged to BTH101 strain bearing the empty plasmids producing T18 and T25 plasmids, where any binding should be considered spurious.

Next, the B2H confirmed candidates underwent a specificity assay. The rationale for the following test is to discard any potential cAMP downstream pathway activation/inhibition beyond the pure TMDs binding. To test the activator specificity, we performed the β-galactosidase assay in the *E. coli* BTH101 strain bearing both plasmids lacking the Pf3::TMD:: 3xGly domains. On the other hand, to test inhibitor specificity, we repeated the enzymatic assay in a BTH101 strain bearing the pUT18-ZIP and pKNT25-ZIP plasmids. These have the cytoplasmic domains in frame fused to the leucine zipper of the GCN4 transcription factor, making the T18-T25 binding highly stable. True activators/inhibitors shall leave the beta-galactosidase activity of these controls unchanged, since any induction or repression in these conditions would mean unspecific stimulation (Figure 2E, 2E’).

We found that 3 out of the 13 selected modulators were specific (dexametasone, parbimostat and varenicline), all of them activators. No inhibitors were found to be specific. Next, we selected the two more promising: parmimostat and varenicline, according to their potential for a chronic treatment. These two drugs where assayed in the human cell line HMC3. Cellular TREM2 was stimulated with agonist Ig for a short time in order not to saturate the Syk -to-Phospho-Syk conversion (p-SYK). This challenge was made in the presence of either DMSO, parbimostat or varenicline at 150 μM. Our results show that varenicline at the assayed dose, increased the p-SYK/SYK ratio ≈40% (Figure 3)

**Figure 3.**
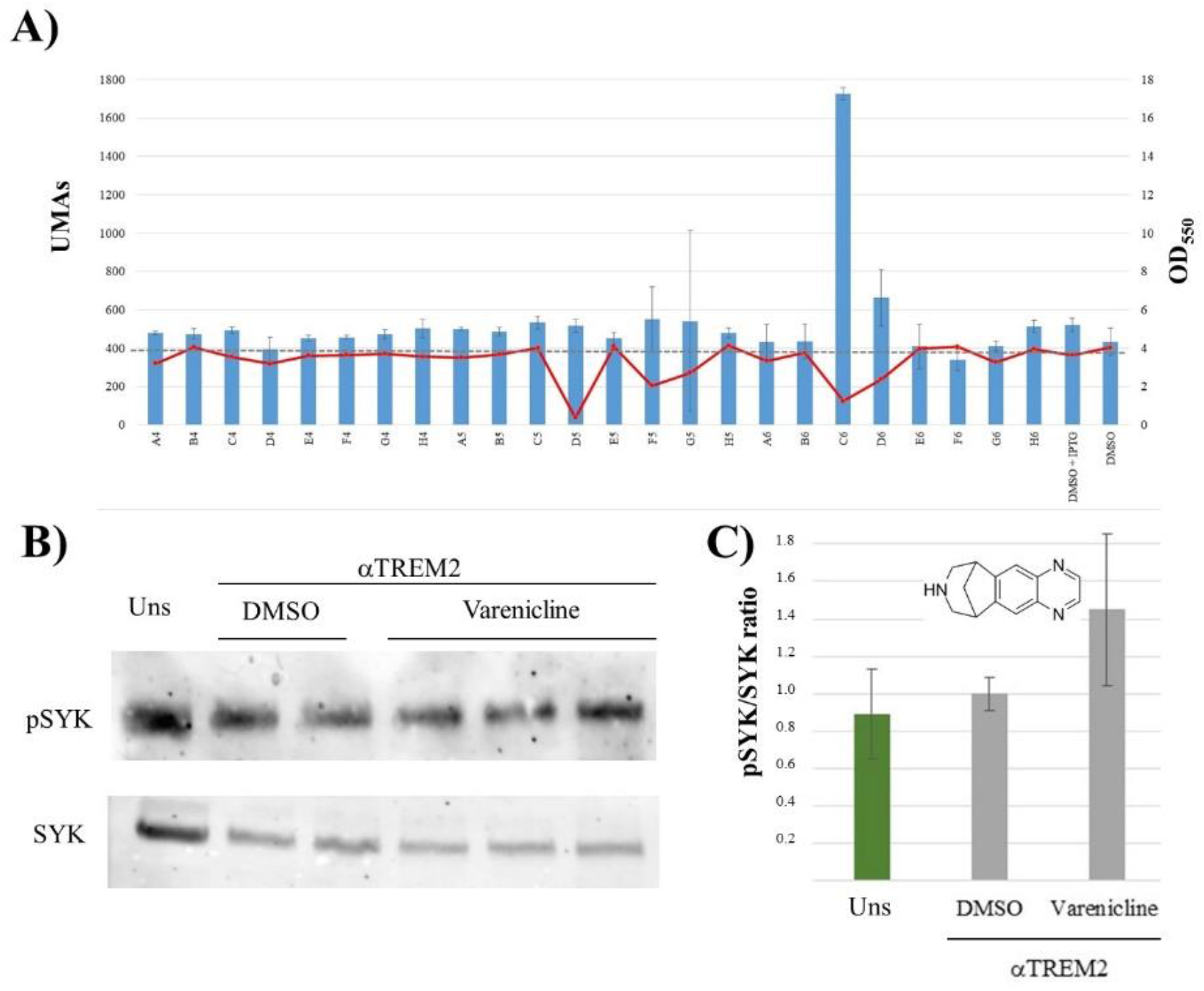
Screening results. A) Representative analysis of the beta-galactosidase activity and bacterial growth *E. coli* BTH101 with the TMD plasmids after O/N stimulation with the different candidates. Candidate in well C6 was selected for further characterization. B) Western blot of HCM3 protein extracts after αTREM2 stimulation with either 1% DMSO or varenicline 150 μM. C) Pathway stimulation after quantifying the different band intensities (n=3).

Varenicline is an organic heterotetracyclic compound that acts as a partial agonist for nicotinic cholinergic receptors currently used as an aid to giving up smoking, however its potential role interacting with TREM2 pathway is completely novel. In order to gain further insight into its effect on the TREM2-TYROBP affinity, we tested their interaction in an *in silico* docking assay. As shown in figure 4B, the best scored binding mode presents the varenicline ligand interacting with both α-helices strands with an affinity value of -5.6 kcal/mol. The non-covalent interactions (NCI) analysis shows that the varenicline ligand is establishing attractive interactions with the ILE-6 and LEU-13 residues from TYROBP’s TMD (right) and with the LEU-6, ALA-7 and PHE-10 residues in chain TREM2’s TMD (left) (Figure 4C).

**Figure 4.**
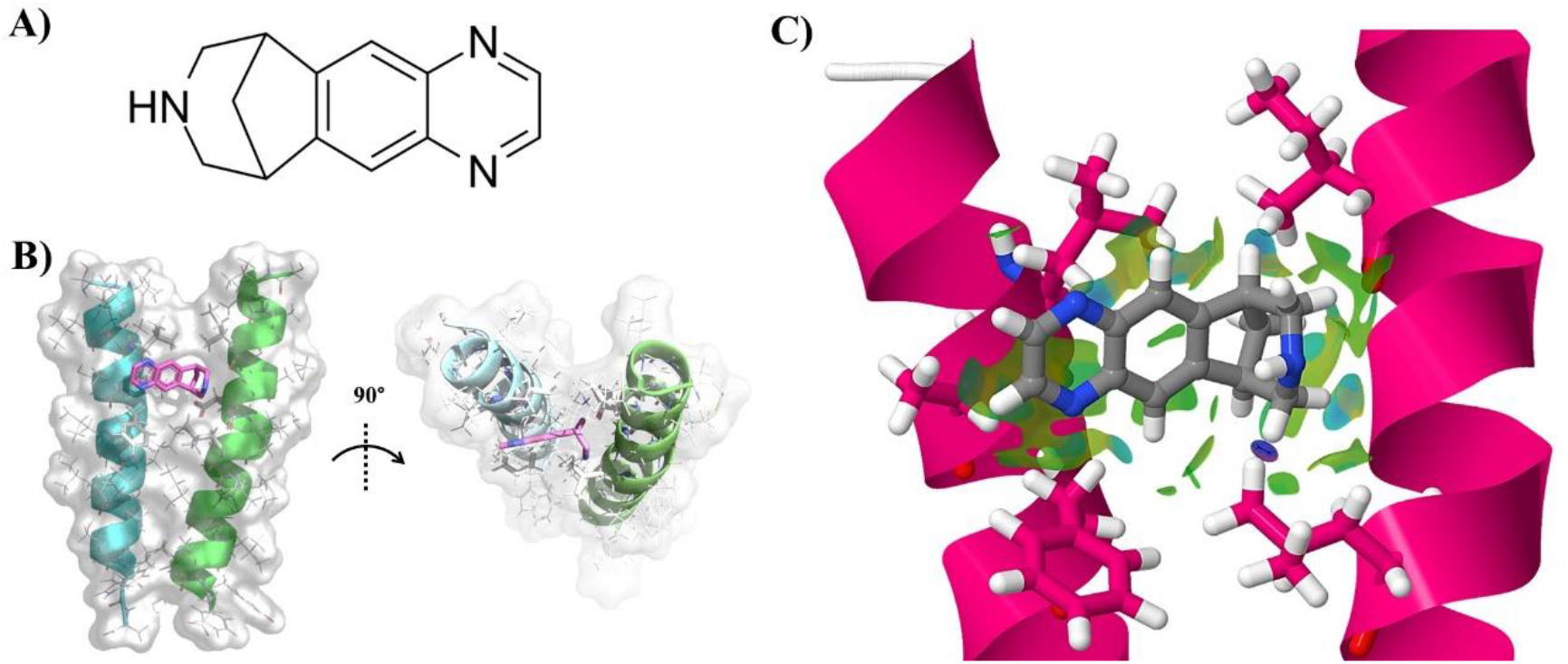
Molecular docking study. A) Varenicline 2D representation. B) Best fitting docking configuration of varenicline with TREM2 (left) and TYROBP (right). C) NCI representation (green area) of the contacts between varenicline and ILE-6 and LEU-13 residues from TYROBP’s TMD (right) and with the LEU-6, ALA-7 and PHE-10 residues in chain TREM2’s TMD (left).

## Discussion

Variants in the TREM2 gene have been strongly associated with an increased risk of AD, highlighting its crucial role in neuroinflammation and microglial response. TREM2 dysfunction contributes to the progression of Alzheimer’s by impairing microglial activity, leading to reduced clearance of amyloid-beta and exacerbated neuronal damage. Activation of TREM2 in early phases of the disease has a positive effect on the evolution of the different mice models of AD (Long et al., 2020; Leng and Edison, 2021; Fracassi et al., 2022). On the other hand, switching off the chronic activation of microglia could impair neurodegeneration and provide a beneficial effect (Deczkowska et al., 2018; Long et al., 2020; Qin et al., 2021). TREM2 has been shown to also interact with other membrane receptors such as DAP10 (Wand S et al 2022). However, it has been demonstrated that the role in AD is closely related with TYROBP (Zhang B et al, 2013). To the best of our knowledge, the current drugs under development specifically target the soluble extracellular domain of TREM2. Here, we postulate that the transmembrane interaction between TREM2 and TYROBP has great potential in pharmacological therapies, given the brain-blood barrier permeability to lipid-soluble small molecules (Pardrige MW, 2012).

The screening of active principles already approved by the FDA offers a wide range of advantages. On one hand, we account with its safety data directly assayed in humans, although these data shall be considered if the drug of choice is required for acute uses, while its eventual use in AD might be considered chronic. In addition, the producing industry already accounts with the necessary permissions in terms of good manufacturing practice, quality controls, etc, what paves the road to a Phase II clinical trial. Repurposing strategies allow development in less time, with lower investment and greater probability of success. Here, we found that varenicline increases TREM2-TYROBP affinity. Varenicline is a nicotine receptor partial agonist currently indicated for smoking cessation. It’s a partial agonist at the α4β2 subtype of the nicotinic acetylcholine might exert a synergic effect with the acetylcholinesterase inhibitors used in AD. Besides, some studies have suggested that varenicline may have neuroprotective effects, meaning it could potentially protect against neurodegeneration. For example, research reported by Bagdas and coworkers (Bagdas D, et al 2018) suggested that varenicline might have neuroprotective effects against Parkinson’s disease-related neurodegeneration in animal models, and administration in aged monkeys improved their cognition and general performance (Terry AV, et al 2016). On the contrary, previous attempts to use varenicline for mild-to-moderate AD failed (<u>NCT00744978</u>), although we cannot discard that any beneficial effect might be relevant if administered in an earlier stage or assayed in a larger cohort (Kim SY, et al 2014). These concerns, however, do not affect the potentiality of our discovery pipeline. Recently, Alector announced results from its INVOKE-2 Phase 2 clinical trial, which evaluated the safety and efficacy of AL002 in people with early AD during 96 weeks. In this trial the clinical diagnosis at enrollment was mild cognitive impairment due to AD for 67% of participants and mild dementia due to AD for 33% of the participants with confirmed brain amyloid accumulation (Alector press release, July 2024). These results underscore the necessity of gaining a more comprehensive understanding of microglial function in AD to efficiently select the right intervention time and target population.

The identification of a potential candidate opens up the possibility of performing a rational design of novel compounds to improve the affinity or pharmacokinetics. Our combination of *in silico* methods and experimental validation, together with further research in the detailed molecular interactions observed in the present study, and its effects on the dimer activity, will allow us to modify the chemical structure of the ligands towards selecting agonist or antagonist effects in the TREM2-TYROBP pathway. We are aware of certain limitations in our pipeline. First, we shall take into account the possibility that gastric or hepatic drug metabolism might change the TREM2-TYROBP binding capacity observed *in vitro*. On the other hand, the native forms of TYROBP in microglia are dimers, while in our assay the interaction assay takes place as monomers. This could imply variations in the interaction and comprise *in vivo* results. Furthermore, promoting interaction might not be directly translated into an intracellular upregulation. It could happen that you simply lengthen the binding time of the complex or eventually reduce the Km between the two domains, causing their affinity to increase. Nevertheless, based on the presented results we are certain that the strategy established in this study will help us overcome these barriers and to obtain promising results.

## Acknowledgements

This project has been funded by Fundacio ACE (Barcelona, Spain) and Fundación SantÁngela (Sevilla, Spain). We also thank Almudena Pino for developing the molecular dynamics and Rubén Laplaza for his helpful comments.

