## Supplementary material for "Drug screening targeting TREM2-TYROBP transmembrane binding"

### Supplementary results

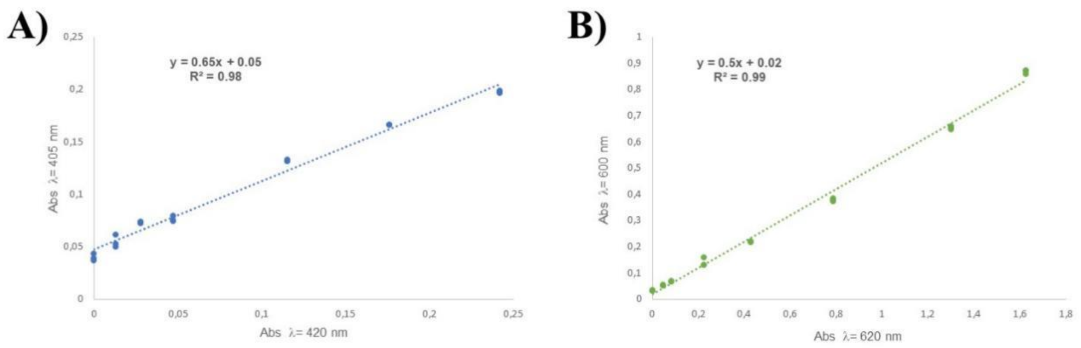


**Supplementary Figure 1. Beta-galactosidase assay absorbance's correlation.** A) Correlation between absorbance at λ = 420 nm (X axis) and λ = 405 nm (Y axis) taken by the ELISA plate reader. Equation and correlation coefficient ($R^2$) can be found in the top left corner. B) Correlation between absorbances at λ = 600 nm (X axis) and λ = 620 nm (Y axis) taken by the ELISA plate reader. Equation and correlation coefficient ($R^2$) can be found in the top left corner. Statistical analysis showed significant results ($p < 0.001$) when performing Spearman's test.

| Antagonists candidates | | | | Agonists candidates | | | |
|---|---|---|---|---|---|---|---|
| **Drug** | **Average** | **StdDev** | **B2H confirmation** | **Drug** | **Average** | **StdDev** | **B2H confirmation** |
| **P163-D9** | 0,2 | 0,03 | Yes | **P161-E6** | 1,16 | 0,03 | - |
| **P163-F6** | 0,5 | 0,05 | Yes | **P161-H6** | 1,17 | 0,07 | - |
| **P162-G11** | 0,5 | 0,10 | - | **P164-G6** | 1,17 | 0,04 | - |
| **P161-A6** | 0,68 | 0,16 | - | **P162-E3** | 1,18 | 0,03 | - |
| **P161-G10** | 0,74 | 0,05 | Yes | **P162-G6** | 1,20 | 0,03 | - |
| **P162-H8** | 0,8 | 0,18 | - | **P162-D4** | 1,25 | 0,07 | - |
| **P162-D5** | 0,76 | 0,05 | - | **P163-C6** | 1,3 | 0,25 | Yes |
| **P162-B8** | 0,8 | 0,10 | - | **P162-G4** | 1,30 | 0,08 | - |
| **P162-H8** | 0,8 | 0,08 | - | **P164-H6** | 1,31 | 0,04 | - |
| **P162-G7** | 0,8 | 0,03 | - | **P161-A5** | 1,33 | 0,14 | Yes |
| **P162-B8** | 0,8 | 0,03 | - | **P164-H5** | 1,36 | 0,15 | - |
| **P164-E9** | 0,90 | 0,14 | - | **P161-E8** | 1,36 | 0,09 | - |
| **P162-D4** | 0,9 | 0,18 | - | **P164-G8** | 1,45 | 0,14 | - |
| | | | - | **P161-F7** | 1,46 | 0,06 | - |
| | | | - | **P164-H8** | 1,49 | 0,08 | - |
| | | | - | **P162-F7** | 1,5 | 0,31 | Yes |
| | | | - | **P164-H4** | 1,50 | 0,14 | - |
| | | | - | **P164-F10** | 1,58 | 0,04 | Yes |
| | | | - | **P164-F6** | 1,59 | 0,04 | - |
| | | | - | **P164-E5** | 1,64 | 0,03 | Yes |
| | | | - | **P164-C6** | 1,66 | 0,15 | Yes |
| | | | - | **P161-H4** | 1,75 | 0,25 | Yes |
| | | | - | **P164-C11** | 2,17 | 0,37 | Yes |
| | | | | **P161-G9** | 2,86 | 0,26 | Yes |
| | | | | **P161-H10** | 3,30 | 1,02 | Yes |

**Supplementary table 1.** Initially selected candidate summary. Induction fold -average and standard deviation- of the different candidate drugs. Basal levels were obtained from bacterial cultures in LB with 1% DMSO (average 1.0 $\pm$ 0.14). As positive control reference, bacterial cultures grown with DMSO plus IPTG was used (average 1.46 $\pm$ 0.72). Only those independently confirmed in an extra beta-galactosidase assay (B2H confirmation column) underwent the specificity assay.

### Supplementary methods

The complete protein used as bait is translated from pKNT25 of the in-frame cloned BATCH pKTN25 plasmid, at the BamHI and HindIII sites, generating the full construct Pf3: TYROBPTMD:

MTMITPSLQSVITDVTGQLTAVQADITTIGGGVLAGIVMGDLVLTVLIALAVYF
LGGGDPRVPSSNSMTMQQSHQAGYANAADRESGIPAAVLDGIKAVAKEKNAT
LMFRLVNPHSTSLIAEGVATKGLGVHAKSSDWGLQAGYIPVNPNLSKLFGRAP
EVIARADNDVNSSLAHGHTAVDLTLSKERLDYLRQAGLVTGMADGVVASNHA
GYEQFEFRVKETSDGRYAVQYRRKGGDDFEAVKVIGNAAGIPLTADIDMFAIM
PHLSNFRDSARSSVTSGDSVTDYLARTRRAAPSI

The full protein translated as prey from the plasmid pUT18  upon an equivalent cloning generated the Pf3: TREM2TMD construct:

MTMITPSLQSVITDVTGQLTAVQADITTIGGSILLLLACIFLIKILAASALWAGGG
DPRVPSSNSAASEATGGLDRERIDLLWKIARAGARSAVGTEARRQFRYDGDMN
IGVITDFELEVRNALNRRAHAVGAQDVVQHGTEQNNPFPEADEKIFVVSATGES
QMLTRGQLKEYIGQQRGEGYVFYENRAYGVAGKSLFDDGLGAAPGVPSGRSK
FSPDVLETVPASPGLRRPSLGAVERQSI
